# MCL-1 inhibition by selective BH3 mimetics disrupts mitochondrial dynamics in iPSC-derived cardiomyocytes

**DOI:** 10.1101/743922

**Authors:** Megan L. Rasmussen, Nilay Taneja, Abigail C. Neininger, Lili Wang, Linzheng Shi, Bjorn C. Knollmann, Dylan T. Burnette, Vivian Gama

## Abstract

MCL-1 is a well characterized inhibitor of cell death that has also been shown to be a regulator of mitochondrial dynamics in human pluripotent stem cells (hPSCs). We used cardiomyocytes derived from hPSCs (hPSC-CMs) to uncover whether MCL-1 is crucial for cardiac function and survival. Inhibition of MCL-1 by BH3 mimetics, resulted in the disruption of mitochondrial morphology and dynamics as well as disorganization of the actin cytoskeleton. Interfering with MCL-1 function affects the homeostatic proximity of DRP-1 and MCL-1 at the outer mitochondrial membrane, resulting in decreased functionality of hPSC-CMs. BH3 mimetics targeting MCL-1 are promising anti-tumor therapeutics. Cardiomyocytes display abnormal functional cardiac performance even after caspase inhibition, supporting a non-apoptotic activity of MCL-1 in hPSC-CMs. Progression towards using BCL-2 family inhibitors, especially targeting MCL-1, depends on understanding not only its canonical function in preventing apoptosis, but also in the maintenance of mitochondrial dynamics and function.

## Introduction

Myeloid cell leukemia-1 (MCL-1) was originally identified as an early-induced gene in human myeloid leukemia cell differentiation (Kozopas et al., 1993; Reynolds et al., 1996; Yang et al., 1996). MCL-1 is structurally similar to other anti-apoptotic BCL-2 (B cell lymphoma-2) family proteins (*i.e.* BCL-2, BCL-XL (B cell lymphoma extra-large)) (Chipuk et al., 2010). However, its larger, unstructured N-terminal domain and shorter half-life likely indicated that MCL-1 was not completely functionally redundant with other anti-apoptotic proteins (Perciavalle and Opferman, 2013). Supporting this idea, MCL-1 has been shown to be essential for embryonic development and for the survival of various cell types, including cardiomyocytes, neurons, and hematopoietic stem cells (Rinkenberger et al., 2000; Thomas et al., 2010; Wang et al., 2013; Opferman, 2016).

MCL-1 is one of the most amplified genes in human cancers and is frequently associated with resistance to chemotherapy (Beroukhim et al., 2010; Perciavalle and Opferman, 2013). Earlier work demonstrated that *MCL-1* genetic deletion is peri-implantation lethal in embryogenesis, not due to defects in apoptosis, but rather due to a combination of an embryonic developmental delay and an implantation defect (Rinkenberger et al., 2000). However, the non-apoptotic mechanism by which MCL-1 functions in normal and cancerous cells is still unclear. We previously reported that MCL-1 regulates mitochondrial dynamics in human pluripotent stem cells (hPSCs, which refers to both human embryonic stem cells (hESCs) and induced pluripotent stem cells (hiPSCS)) (Rasmussen et al., 2018). We found that MCL-1 maintains mitochondrial network homeostasis in hPSCs through interactions with Dynamin related protein-1 (DRP-1) and Optic atrophy type 1 (OPA1). In this study, we investigated whether this non-apoptotic role of MCL-1 remains as stem cells differentiate, using cardiomyocytes derived from human induced pluripotent stem cells (hiPSC-CMs).

Mitochondrial fusion promotes elongation of the mitochondrial network, which is key for mitochondrial DNA (mtDNA) homogenization and efficient assembly of the electron transport chain (ETC) (Westermann, 2010; Friedman and Nunnari, 2014). Loss of mitochondrial fusion has been implicated as a mechanism for the onset of dilated cardiomyopathy (Dorn, 2013). Mitochondria also regulate cardiomyocyte differentiation and embryonic cardiac development (Kasahara et al., 2013; Kasahara and Scorrano, 2014; Cho et al., 2014). However, there is limited information about the mechanisms used by cardiomyocytes to minimize the risks for apoptosis, especially in cells derived from highly sensitive stem cells (Imahashi et al., 2004; Murriel et al., 2004; Gama and Deshmukh, 2012; Dumitru et al., 2012; Walensky, 2012).

Ultrastructural changes have long been observed in response to alterations in oxidative metabolism (Hackenbrock, 1966; Khacho et al., 2016). It has become increasingly clear that individual mitochondrial shape changes can also have dramatic effects on cellular metabolism. Mitochondrial morphology and cristae structure are influenced by fission and fusion events; subsequently, ETC complexes that reside on the inner mitochondrial membrane are disrupted upon aberrant fission (Chan, 2007). Several human diseases, such as MELAS (Muscle atrophy, Encephalopathy, Lactic Acidosis, Stroke-like episodes) and Dominant Optic Atrophy (DOA), are associated with mutations in the mitochondrial dynamics and mitochondrial metabolism machineries (Chan, 2007; Hsu et al., 2016). Likewise, many neurological conditions, including Parkinson’s disease, Huntington’s disease, and Charcot-Marie Tooth Type 2 syndrome, can originate from alterations in mitochondrial dynamics and metabolic regulation (Itoh et al., 2013; Burté et al., 2015). Besides neurological conditions, several studies in the heart suggest that alterations in mitochondrial dynamics causes abnormal mitochondrial quality control, resulting in the buildup of defective mitochondria and reactive oxygen species (ROS) (Galloway and Yoon, 2015; Song et al., 2017). Interestingly, it has been shown that modulating the production of ROS can favor or prevent differentiation into cardiomyocytes (Buggisch et al., 2007; Murray et al., 2014). Thus, specific metabolic profiles controlled by mitochondrial dynamics are likely critical for hiPSC-CMs, since they can influence cell cycle, biomass, metabolite levels, and redox state (Zhang et al., 2012).

It is not completely understood how dynamic changes in metabolism affect cardiomyocyte function. Deletion of MCL-1 in murine heart muscle resulted in lethal cardiomyopathy, reduction of mitochondrial DNA (mtDNA), and mitochondrial dysfunction (Wang et al., 2013). Inhibiting apoptosis via concurrent BAK/BAX knockout allowed for the survival of the mice; conversely, the mitochondrial ultrastructure abnormalities and respiratory deficiencies were not rescued. These results indicate that MCL-1 also has a crucial function in maintaining cell viability and metabolic profile in cardiomyocytes. Despite these efforts, the non-apoptotic mechanism by which MCL-1 specifically functions in cardiomyocytes is still unknown. Furthermore, a role for MCL-1 in the regulation of mitochondrial dynamics in cardiac cells has not yet been defined. Here we describe findings that MCL-1 is essential for the survival of hiPSC-CMs by maintaining mitochondrial morphology and function.

## Results and Discussion

### MCL-1 inhibition causes severe defects in hiPSC-CM mitochondrial network

Recently published small molecule inhibitors of MCL-1 have been anticipated as potent anti-tumor agents against MCL-1-dependent cancers with limited cardiotoxicity in mouse models (Cohen et al., 2012; Kotschy et al., 2016; Letai, 2016). Thus, we chose to use hiPSC-CMs (Figure 1A) to examine the effects of MCL-1 inhibition on mitochondrial morphology, using the small molecule inhibitor S63845 (Kotschy et al., 2016), combined with structured illumination microscopy (SIM) to observe mitochondria at high-resolution (Figure 1B). Cardiomyocytes were images after 4 days of treatment with vehicle (DMSO) or MCL-1 inhibitor (MCL-1*i*/S63845) and the caspase inhibitor Q-VD-OPh (QVD) (Figure 1B). We found that MCL-1 inhibition had significant effects on iPSC-CM mitochondrial morphology. Mitochondrial networks in S63845-treated cells were severely disrupted, with individual mitochondria becoming more fragmented and globular, as opposed to elongated and interconnected networks in control cells (Figure 1C). In a previous report, MCL-1 inhibition using RNAi also resulted in mitochondria morphology defects including severe cristae disruption and remarkable vacuolation in the mitochondrial matrix (Guo et al., 2018). Recent reports have determined that MCL-1 functions not only as an apoptosis regulator but also as a modulator of mitochondrial morphology and dynamics (Perciavalle et al., 2012; Morciano et al., 2016; Rasmussen et al., 2018). Thus, we hypothesized that inhibiting MCL-1 with BH3 mimetics would affect the functionality of human cardiomyocytes, due to the disruption of crucial MCL-1 interactions with the mitochondrial dynamics machinery, which ultimately will lead to cell death.

**Figure 1:**
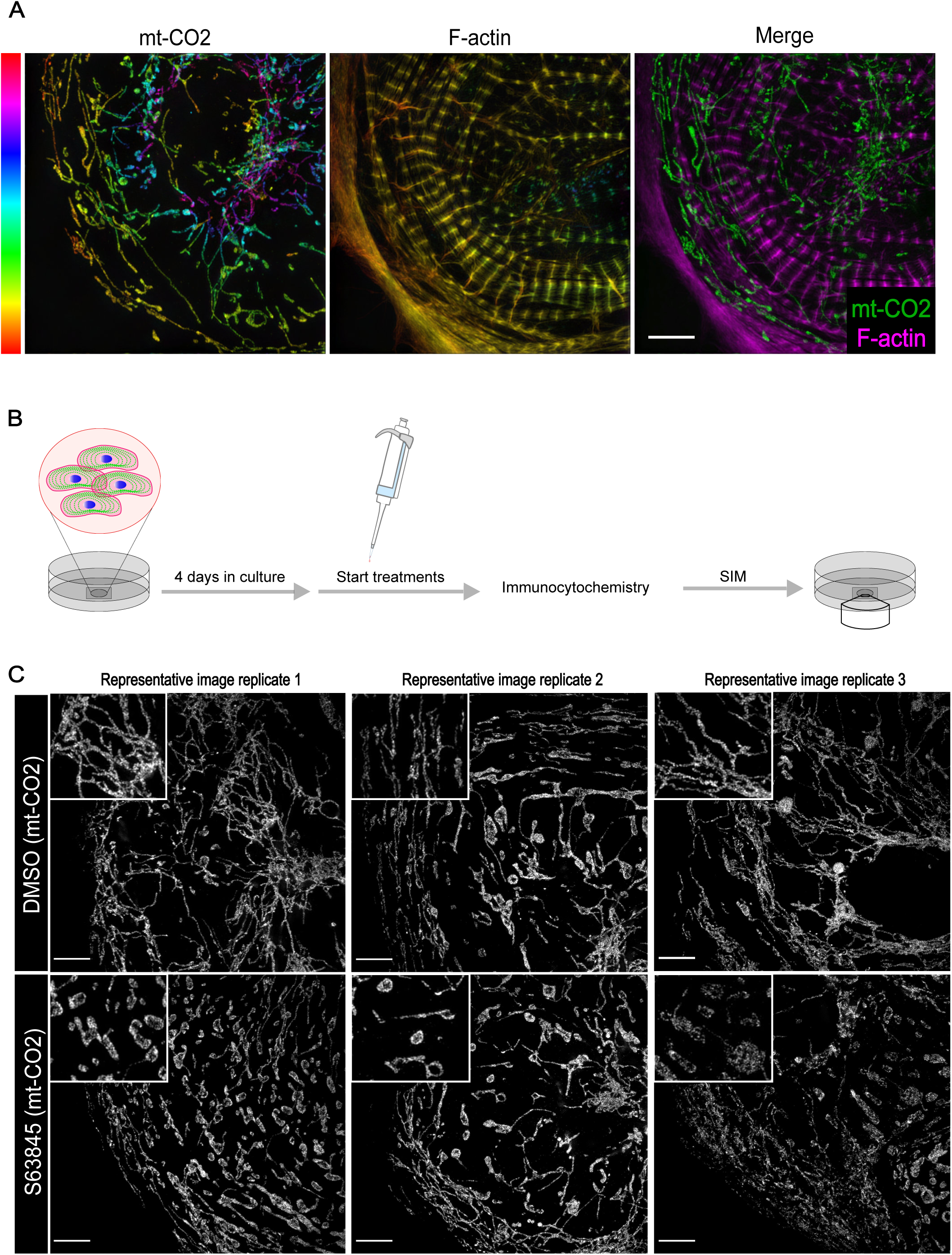
MCL-1 inhibition causes mitochondrial fragmentation. (A) Maximum intensity projection showing mitochondria (mt-CO2) and myofibril (F-actin) organization in an untreated hiPSC-CM. Rainbow LUT shown to denote Z-depth. Scale: 10 μm. (B) Schematic of cell treatment paradigm used throughout this study. Structured Illumination Microscopy (SIM) was used for acquisition of all super-resolution images. (C) hiPSC-CMs were treated with vehicle (DMSO) or 2 μM S63845 and Q-VD-Oph (QVD). Vehicle-treated cells have elongated mitochondria assembled in networks; MCL-1 inhibition causes mitochondria to become fragmented and disorganized. Insets show magnification of individual mitochondria morphology. Scale: 10 μm. Representative images are shown for all panels.

### MCL-1 inhibition affects contractility of iPSC-CMs and myofibril assembly in a caspase-independent manner

MCL-1 inhibition by S63845 was shown to have minimal effects on murine ejection fraction (Kotschy et al., 2016) and on overall cardiac function in human cardiomyocytes (Guo et al., 2018). These results are intriguing considering previous studies reporting that MCL-1 deletion from murine cardiomyocytes has severe effects on mitochondrial morphology and cardiac function, which were not rescued by co-deletion of BAK and BAX (Wang et al., 2013). We treated human iPSC-CMs with S63845, while inhibiting caspase activity, and measured spontaneous beating using phase-contrast live-cell imaging. We observed lower numbers of beating cells when treated with 1-2 μM MCL-1*i* (S63845), and the cells that were beating appeared to beat more slowly (Figure S1A-C). To assess these defects more rigorously, we plated cells on a multi-electrode array (MEA) plate and examined cardiac function using the Axion Biosystems analyzer (Clements and Thomas, 2014) (Figure 2A). We observed that MCL-1 inhibition caused severe defects in cardiomyocyte functionality after just 18 hours of the first treatment (Figure 2B-D). In particular, beat period irregularity was significantly increased (Figure 2B), while spike amplitude and spike slope means were decreased (Figure 2C-D). The differences between beat period mean and conduction velocity mean at this time point were not significant (Figure S1D-E); however, at just two days of treatment with MCL-1 inhibitor, cardiomyocytes became quiescent and stopped beating altogether (Figure S1F-J). These results implicate tachycardia and arrhythmia phenotypes in cardiomyocytes exposed to S63845. To probe whether these cells are also sensitive to BCL-2 inhibition, we also treated hiPSC-CMs with the BCL-2 inhibitor Venetoclax (ABT-199) (Souers et al., 2013). In the same treatment paradigm, ABT-199 had no effect on hiPSC-CM functionality compared to control cells (Figure S1C-G). These results suggest that hiPSC-CMs are highly dependent on MCL-1, but not BCL-2, for function and survival. Intriguingly, we also observed significant changes in the structure and integrity of the actin network and subsequent myofibril organization in cells treated with MCL-1 inhibitor (Figure 2E). hiPSC-CMs displayed poor Z-line organization, lower density of F-actin, and increased presence of stress fibers (Figure 2E). Blinded quantification of F-actin organization revealed that MCL-1 inhibitor-treated cells had significantly less organized myofibril structure (Figure 2F).

**Figure 2:**
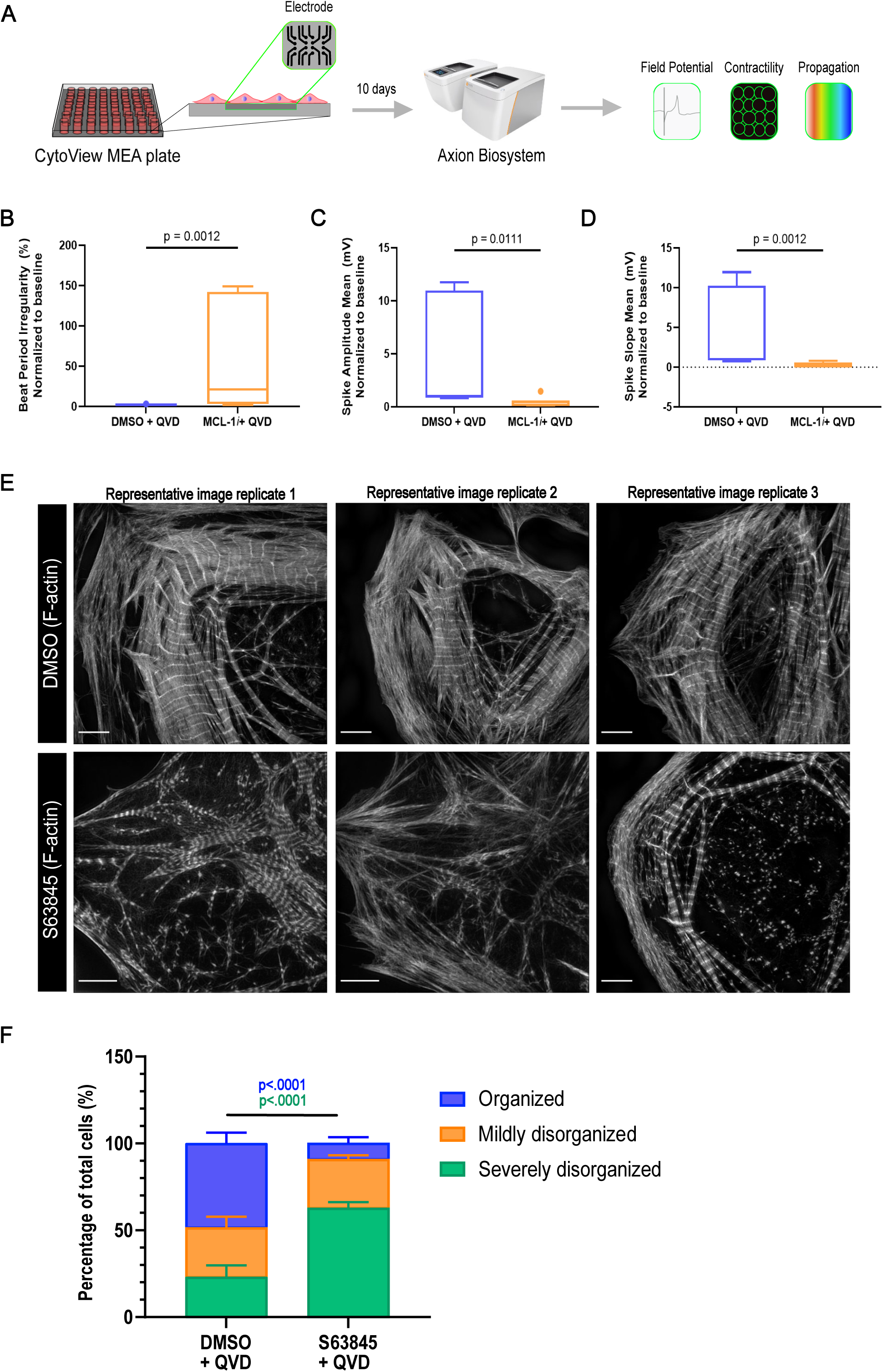
MCL-1 inhibition causes disruption of myofibrils and functional defects. (A) Schematic of Axion Biosystems MEA paradigm for recording cardiac performance in live cells. hiPSC-CMs were plated on a CytoView MEA plate (Axion Biosystems) and treated with either vehicle (DMSO) or .5 μM MCL-1*i* (S63845) and QVD. Live-cell activity was recorded at 18 hours-post-treatment for 5 minutes; (B) beat period irregularity was increased in MCL-1*i*-treated cells, while spike amplitude mean and spike slope mean were decreased (C-D). (E) Vehicle-treated hiPSC-CMs have organized myofibril structure as shown by maximum intensity projections. hiPSC-CMs treated with 2 μM MCL-1*i* (S63845) and QVD have myofibrils that are unorganized and poorly defined Z-lines. Scale: 10 μm. Representative images are shown for all panels. (F) Quantification of myofibril structure phenotypes represented in Figure 2E (n=~80 cells from 3 separate experiments).

### MCL-1 co-localizes with mitochondrial dynamics proteins in hiPSC-CMs, and S63845 disrupts MCL-1:DRP-1 co-localization

Since MCL-1 inhibition disrupted mitochondrial network integrity in hiPSC-CMs and MCL-1 depletion affects mitochondrial dynamics proteins (Rasmussen et al., 2018), we next examined the effects of MCL-1 inhibition on the expression levels of key mitochondrial proteins. MCL-1 inhibitor-treated cells had a significant increase in the expression levels of DRP-1 (Figure 3A-B) and MCL-1 (Figure 2C-D). Previous studies using S63845 (Kotschy et al., 2016) also reported the induction of MCL-1 expression. There were no significant changes in the expression levels of phospho-DRP-1 (pDRP-1 S616), OPA1 or TOM20 (Figure 2C-D and Figure S3A). We then assessed whether MCL-1 interacts with these GTPases responsible for maintaining mitochondrial morphology and dynamics using *in situ* proximity ligation assay (PLA). Our data shows that MCL-1 is in close proximity to both DRP-1 and OPA1 (Figure 3E-H). PLA puncta were quantified and normalized to the number of puncta in the control sample (Figure S2B). The co-localization of MCL-1 with DRP-1, but not OPA1, was disrupted upon inhibition of MCL-1 with S63845 (Figure 3E-F), suggesting that MCL-1 interacts with DRP-1 through its BH3 binding groove. Since the interaction with OPA1 was not disturbed (Figure 3G-H), it is possible that MCL-1 interacts with OPA1 either through a different domain, or with a different isoform of OPA1 in hiPSC-CMs than in hPSCs (Rasmussen et al., 2018). Another possibility is that, upon differentiation, the small molecule can no longer penetrate the inner mitochondrial membrane.

**Figure 3:**
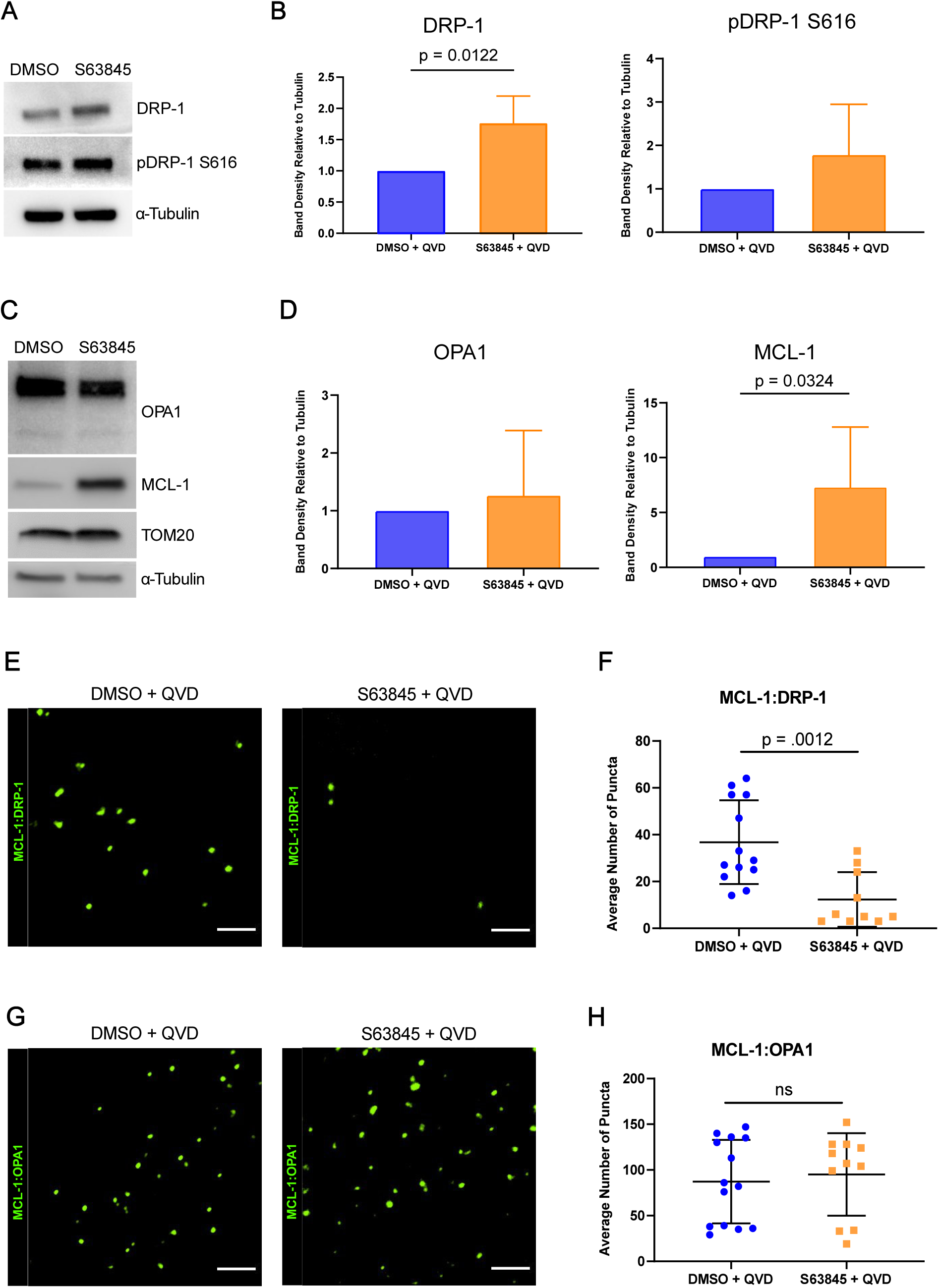
MCL-1 interacts with mitochondrial dynamics proteins. (A) Western blot showing DRP-1 activity in hiPSC-CMs treated with S63845 + QVD. (B) Quantification of DRP-1 and pDRP-1 S616 band density relative to α-tubulin. (C) Western blot showing OPA1, MCL-1, and TOM20 levels in hiPSC-CMs treated with S63845 + QVD. (D) Quantification of OPA1 and MCL-1 band density relative to α-tubulin. Images from PLA showing representative ROIs showing MCL-1:DRP-1 (E) or MCL-1:OPA1 (G) puncta in vehicle- or S63845-treated hiPSC-CMs (Scale: 5 μm). Quantification of PLA puncta from MCL-1:DRP-1 (F) or MCL-1:OPA1 (H) interactions (n = 10-15 images per condition from 3 independent experiments). All error bars indicate ±SD.

DRP-1 is shuttled to the outer mitochondrial membrane upon activation. In our previous study, we showed that MCL-1 depletion decreases the activity of DRP-1 and promotes mitochondrial elongation (Rasmussen et al., 2018). Since MCL-1 inhibition with S63845 appeared to cause mitochondrial fragmentation in cardiomyocytes, we hypothesized that more DRP-1 would be activated and brought to the mitochondria to initiate fission. However, levels of active DRP-1 (pDRP-1 S616) that co-localized with mitochondria decreased in S63845-treated hiPSC-CMs (Figure S3A-B). To further assess the disruption of the mitochondrial network caused by MCL-1 inhibition, we employed an assay using a photo-convertible plasmid (mito-tdEos) to assess connectivity and fusion/motility of mitochondria. After photo-conversion, we saw that both the initial converted area and the spread of the converted signal after 20 minutes were significantly decreased (Figure 4A-D). This fragmentation caused by MCL-1 inhibition was also DRP-1 dependent, since knockdown of DRP-1 rescued the increased fragmentation in S63845-treated cells (Figure 4E-F and Figure S3C). The recruitment of DRP-1 to the mitochondria has been proposed to be a critical inducer of mitophagy (Lee et al., 2011; Kageyama et al., 2014; Burman et al., 2017). Thus, an interesting possibility is that inhibition of MCL-1 is decreasing clearing of damaged mitochondria in cardiomyocytes due to the decrease in recruitment of active DRP-1.

**Figure 4:**
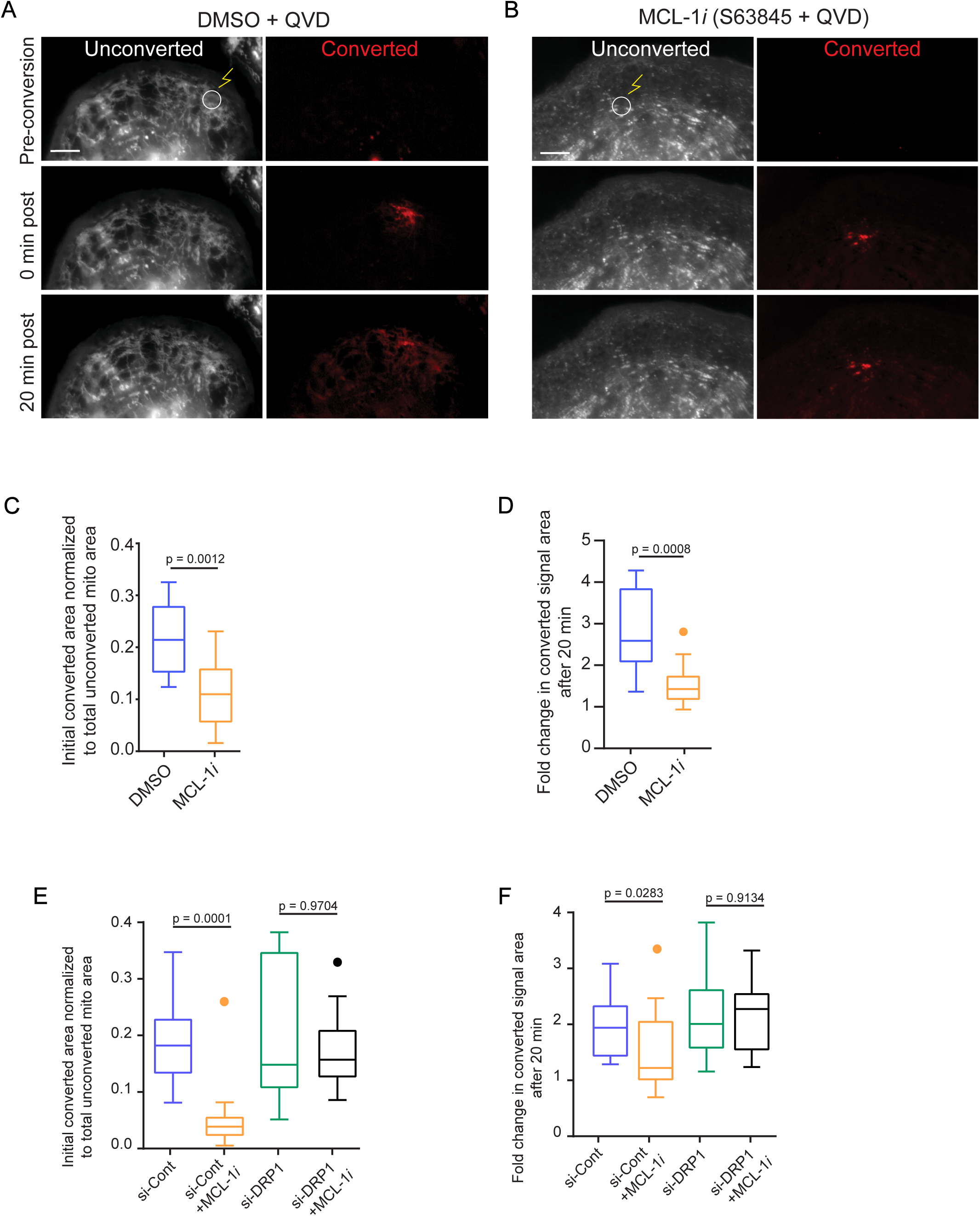
MCL-1 inhibition results in mitochondrial fragmentation through a DRP-1 dependent manner. (A) Vehicle- and (B) S63845-treated hiPSC-CMs were transfected with Mito-tdEos and a small area was photo-converted (see methods). Cells were imaged for 20 minutes-post-conversion to assess mitochondrial network connectivity. Quantification of (C) initial converted area normalized to total unconverted area and (D) fold change in converted area after 20 minutes from Figure 4A-B. (E) Quantification of initial converted area normalized to total unconverted area in hiPSC-CMs treated with si-Control (si-Cont) ±MCL-1*i* (2 μM) and si-DRP-1 ±MCL-1*i* (2 μM). (F) Quantification of fold change in converted area after 20 minutes in same treatments from Figure 4E. Boxplots show Tukey whiskers.

### MCL-1 inhibition results in iPSC-CM death

To examine whether iPSC-CMs treated with MCL-1 inhibitor were still sensitive to caspase-mediated cell death, we treated the cells with increasing doses of S63845 and examined the activation of caspase-3 and caspase-7 in the absence of QVD. Cells responded to S63845 in a dose-dependent manner after 48 hours, with 1-2 μM MCL-1*i* inducing the most caspase activity (Figure 5A). To examine the possibility that cardiomyocytes were dying independently of caspase-3 activation, we used the Incucyte live cell imaging system and indeed found similar levels of cell death with and without caspase inhibition (Figure S4A-B). These results indicate that iPSC-CMs are committing to a caspase-independent cell death in response to MCL-1 inhibition. Previous reports have established that iPSC-derived cardiomyocytes are more similar to immature progenitor cells. To test the possibility that the effects were caused by this immature state, we used a previously published hormone-based method for cardiomyocyte maturation (Figure 5B-C) (Parikh et al., 2017). We tested for caspase-3/7 activation after 24 hours of treatment with increasing doses of S63845 and detected similar effects of MCL-1 inhibition in hormone-matured hiPSC-CMs and vehicle-treated hiPSC-CMs. (Figure 5D-E). These results together with previous work from other groups (Thomas et al., 2013; Wang et al., 2013) highlight the importance of extended and rigorous testing of safety and potential off-target effects of MCL-1 inhibitors on human cardiomyocytes.

**Figure 5:**
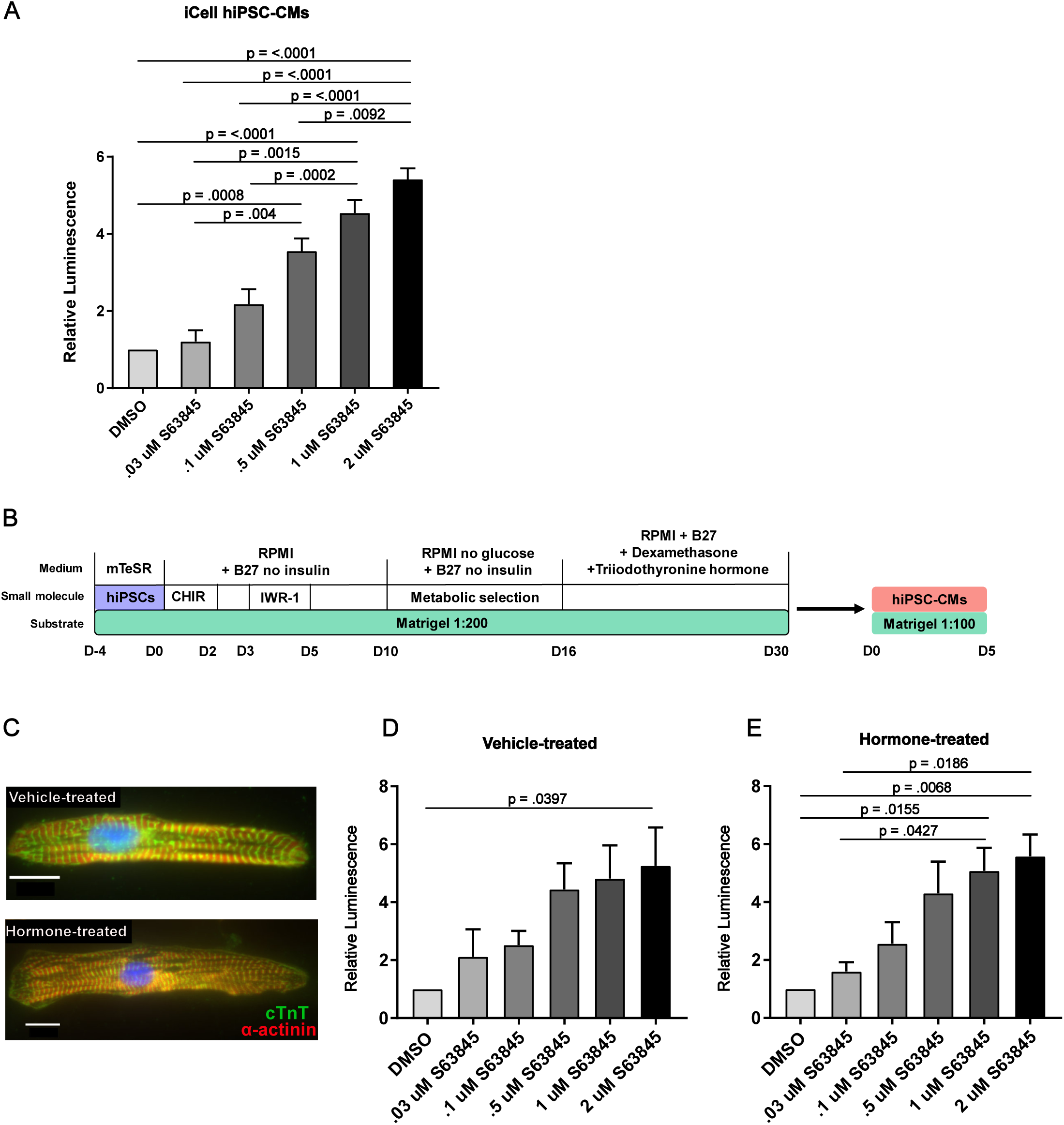
hiPSC-CMs commit to intrinsic apoptosis after MCL-1 inhibition. (A) iCell hiPSC-CMs were treated with increasing doses of S63845 for 48 hours before caspase activity was measured by CaspaseGlo 3/7 assay (Promega). (B) Schematic of maturation protocol for hiPSC-CMs shown in Figure 1C. (C) hiPSC-CMs treated with Dex (dexamethasone) and T3 (triiodothyronine) display more mature phenotype compared to vehicle-treated control cells. (D) Vehicle- or (E) Dex+T3-treated hiPSC-CMs were exposed to S63845 at increasing doses for 24 hours. Caspase activity was measured as in Figure 1A.

### Long-term MCL-1 inhibition, but not BCL-2 inhibition, causes defects in cardiomyocyte functionality

MCL-1 inhibition has significant effects on hiPSC-CM contractility and functionality when used at higher doses (Figure 2B-D and Figure S1C-G). To test if MCL-1 inhibition still depletes cardiac functionality at lower doses, we treated hiPSC-CMs for two weeks (with treatments every two days) with 100 nM S63845. We also treated the cells with the BCL-2 inhibitor ABT-199 (100 nM) and a combination of S63845 + ABT-199 (100 nM each). While there were no significant differences between treatments in the spike slope mean (Figure 6B) and beat period mean (Figure S5A), either MCL-1 inhibition alone or the combination treatment significantly disrupted hiPSC-CM spike amplitude mean (Figure 6A), conduction velocity mean (Figure 6C), max delay mean (Figure 6D), propagation consistency (Figure 6E), and field potential duration (Figure S5B). Cells treated with ABT-199 appeared healthy and were functionally similar to control cells throughout the experiment (Figure 6A-E and Figure S5A-B). Cells displayed mitochondrial network and actin disruption in the S63845-treated condition, and even more severe phenotypes were observed in cells treated with both inhibitors when compared to control cells (Figure 6F-I and Figure S5C-F). BCL-2 inhibition had little effect on mitochondrial network organization and virtually no effect on myofibril organization (Figure 6H and Figure S5E). These results further support the idea that MCL-1 plays an important role in the mitochondrial homeostasis of cardiomyocytes. It would be of interest to determine whether MCL-1 function in mitochondrial dynamics affects the maturation of iPSC-CMs or heart development *in vivo* (Kasahara et al., 2013; Feaster et al., 2015; Parikh et al., 2017). We speculate that other determinants of mitochondrial homeostasis, including mitochondrial biogenesis and mitophagy, may be affected by MCL-1 deficiency in these cells as they mature.

**Figure 6:**
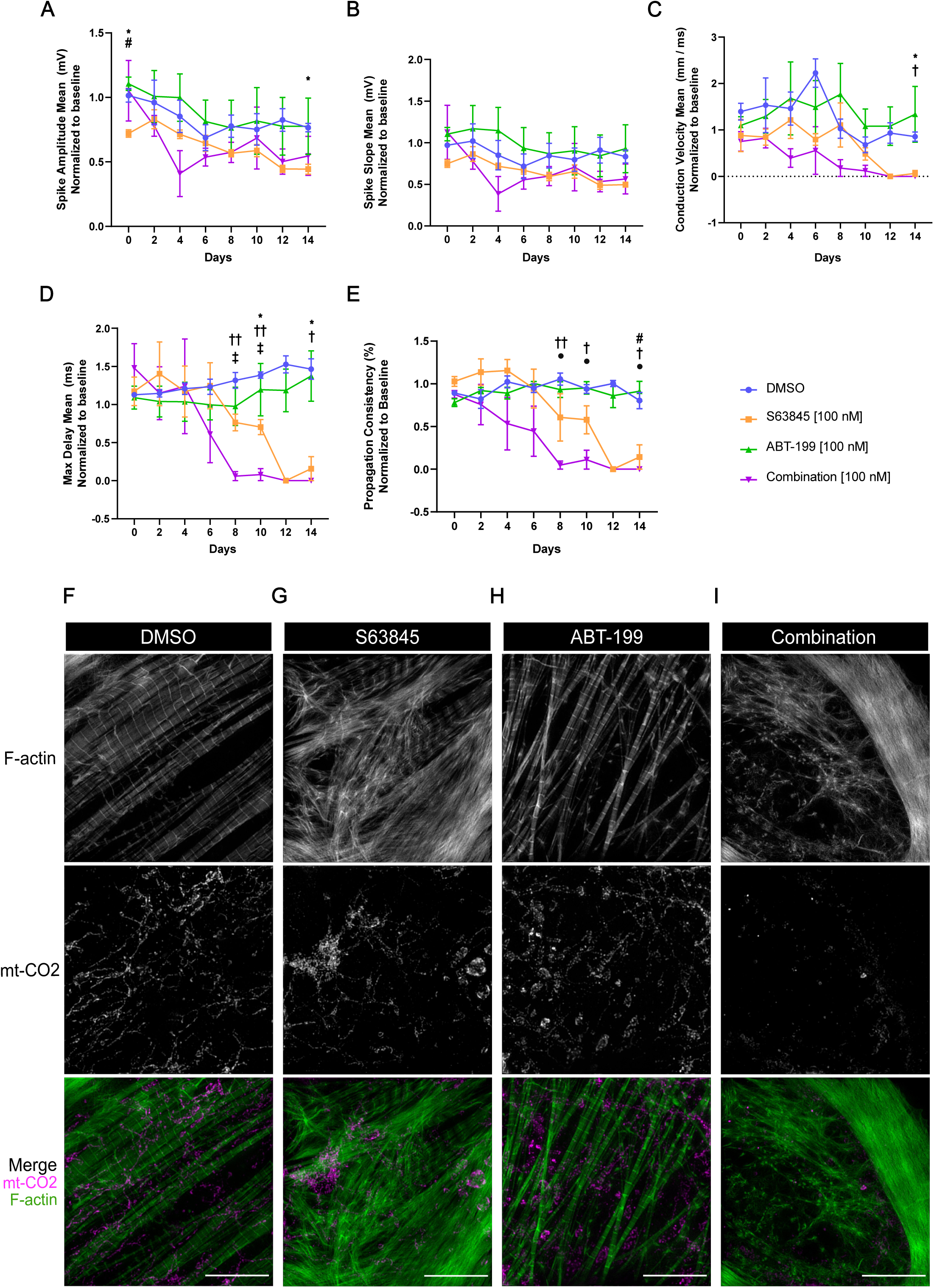
Chronic inhibition of MCL-1, but not BCL-2, results in cardiac activity defects. hiPSC-CMs were treated every 2 days with DMSO (blue), 100 nM S63845 (orange), 100 nM ABT-199 (green), or both inhibitors (magenta) for 14 days. MEA plate was recorded 2 hours-post-treatment for 5 minutes and results were normalized to baseline recording for each respective well. Results of recordings for spike amplitude mean (A), spike slope mean (B), conduction velocity mean (C), max delay mean (D), and propagation consistency (E) are shown. P-values show significance as follows: * = DMSO vs. S63845, † = DMSO vs. Combination, # = S63845 vs. ABT-199, ‡ = S63845 vs. Combination, ● = ABT-199 vs. Combination. One symbol indicates p = <0.05, two symbols indicate p = <0.01. Error bars indicate ±SEM. (F-I) Mitochondria and F-actin were imaged at the end of the treatment paradigm in Figure 6A-E. Representative images are shown of cells treated with DMSO (F), 100 nM MCL-1*i* (S63845) (G), 100 nM BCL-2*i* (ABT-199) (H), and 100 nM MCL-1*i* + 100 nM BCL-2*i* (Combination) (I). Scale: 10 μm.

Whether the function of MCL-1 in mitochondrial dynamics is critical for maintaining the metabolic profile of iPSC-CMs is not known. Studies from our laboratory show that inhibition of MCL-1 induces the differentiation of iPSCs (Rasmussen et al., 2018), which is likely associated with changes in metabolism to support cell-type specific processes (Folmes et al., 2016). Since mitochondrial morphology is tightly coupled to cellular respiration via integrity of the ETC, future studies will aim to investigate the metabolic changes that occur when MCL-1 is deleted in iPSC-CMs. Cardiac contractions depend on energy from these metabolic pathways, and thus cardiac mitochondria are forced to work constantly and likely require strict quality control mechanisms to maintain a functioning state (Dorn et al., 2015). This quality control process could depend in part on MCL-1. In support of this idea, our studies indicate that MCL-1 activity is essential for iPSC-CM viability and maturation, which could be linked to MCL-1’s non-apoptotic function at the mitochondrial matrix. Our results emphasize the need for a more complete molecular understanding of MCL-1’s mechanism of action in human cardiomyocytes as it may reveal new approaches to prevent potential cardiac toxicities associated with chemotherapeutic inhibition of MCL-1.

## Materials and Methods

### Cell Culture

Human induced pluripotent stem cell-derived cardiomyocytes (iCell Cardiomyocytes^2^) were obtained from Cellular Dynamics International (#CMC-100-012-000.5). Cells were thawed according the manufacturer protocol in iCell Plating medium. Briefly, cells were thawed and plated on 0.1% gelatin at 50,000 cells/well in 96-well plates. Cells were maintained at 37°C and 5% CO_2_ and fed every other day with iCell Cardiomyocyte Maintenance medium (Cellular Dynamics International #M1003). For knockdown experiments, wells were coated with 5 μg/mL fibronectin (Corning #354008) 1 hour prior to plating. For functional experiments using the Axion bioanalyzer, cells were plated on 50 μg/mL fibronectin in a 48-well CytoView MEA plate (Axion Biosystems #M768-tMEA-48B). For imaging experiments, cells were re-plated on glass-bottom 35 mm dishes (Cellvis #D35C4-20-1.5-N) coated with 10 μg/mL fibronectin. For live-cell imaging, cells were maintained at 37°C with 5% CO^2^ in a stage top incubator (Tokai Hit).

### Cell Treatments

All treatments were added directly to cells in iCell Cardiomyocyte Maintenance media. The pan-caspase inhibitor Q-VD-OPh (SM Biochemicals #SMPH001) was added to cells at a concentration of 25 µM. The small molecule MCL-1 inhibitor derivative (S63845) was a gift from Joseph Opferman (St. Jude’s Children Hospital). ABT-199 was purchased from Active Biochemicals (#A-1231). All stock solutions were prepared in DMSO.

### RNAi and Plasmid Transfection

Commercially available siRNA targeting DRP-1 (Thermo Fisher Scientific # AM51331) was used to generate transient knockdowns in hiPSC-CMs. Cells were seeded at 50,000 cells per well in a 96-well plate coated with 5 μg/mL fibronectin. Cells were transfected as per the manufacturer protocol using TransIT-TKO Transfection Reagent (Mirus Bio #MIR2154) in iCell maintenance media containing 25uM Q-VD-OPh. To increase knockdown efficiency, the transfection was repeated 48 hours later. Cells were left to recover for an additional 24 hours in fresh media containing 25uM Q-VD-OPh. Cells were lysed for Western blot or re-plated on glass-bottom 35 mm dishes and fixed for analysis by immunofluorescence. Silencer Select Negative Control No. 1 (Thermo Fisher Scientific # 4390843) was used as a control.

Plasmid encoding mito-tdEos (Addgene #57644) was transfected using ViaFect (Promega #E4981) as described in the manufacturer protocol. Cells were maintained until optimal transfection efficiency was reached before cells were imaged.

### Immunofluorescence

For immunofluorescence, cells were fixed with 4% paraformaldehyde for 20 min and permeabilized in 1% Triton-X-100 for 10 min at room temperature. After blocking in 10% BSA, cells were treated with primary and secondary antibodies using standard methods. Cells were mounted in Vectashield (Vector Laboratories #H-1000) prior to imaging. Primary antibodies used include Alexa Fluor-488 Phalloidin (Thermo Fisher Scientific #A12379), mouse anti-mtCO2 (Abcam #ab110258), rabbit anti-pDRP-1 S616 (Cell Signaling Technologies #3455S). For Incucyte experiments, nuclei were visualized using NucLight Rapid Red Reagent (Essen Bioscience #4717). Alexa Fluor-488 (Thermo Fisher Scientific #A11008) and Alexa Fluor-568 (Thermo Fisher Scientific #A11011) were used as secondary antibodies. MitoTracker Red CMXRos (Thermo Fisher Scientific #M7512) added at 100 nM was used to visualize mitochondria in PLA experiments.

### Western blot

Gel samples were prepared by mixing cell lysates with LDS sample buffer (Life Technologies, #NP0007) and 2-Mercaptoethanol (BioRad #1610710) and boiled at 95°C for 5 minutes. Samples were resolved on 4-20% Mini-PROTEAN TGX precast gels (BioRad #4561096) and transferred onto PVDF membrane (BioRad #1620177). Antibodies used for Western blotting are as follows: DRP-1 (Cell Signaling Technologies #8570S), pDRP-1 S616 (Cell Signaling Technologies #4494), OPA1 (Cell Signaling Technologies #67589S), MCL-1 (Cell Signaling Technologies #94296S), TOM20 (Cell Signaling Technologies # 42406S), and α-Tubulin (Sigma # 05-829).

### Impedance assays

The Axion Biosystems analyzer was used to measure contractility and impedance in iPSC-CMs. Cells were plated on 48-well CytoView MEA plates and maintained for 10 days before treatment and recordings. Recordings were taken for 5 minutes approximately two hours after media change. Cells were assayed using the standard cardiac analog mode setting with 12.5 kHz sampling frequency to measure spontaneous cardiac beating. The Axion instrument was controlled using Maestro Pro firmware version 1.5.3.4. Cardiac beat detector settings are as follows:

|  |  |
| --- | --- |
| Beat Detection Threshold | 300 μV |
| Min. Beat Period | 250 ms |
| Max. Beat Period | 5 s |
| Synchronized Beat Maximum Propagation Delay | 30 ms |
| Minimum Active Channels Ratio | 50.00% |
| Running Average Beat Count | 10 |

### Proximity Ligation Assay (PLA)

Cells were seeded onto fibronectin-coated 8-chamber MatTek glass slides (#CCS-8) at 10,000 cells/chamber. After treatments, cells were fixed in 4% PFA for 20 min and permeabilized in 1% Triton-100-X for 10 min at room temperature. Following fixation, the DuoLink proximity ligation assay (Sigma #DUO92014) was performed as per manufacturer protocol. The primary antibodies were incubated overnight at 4°C and are as follows: mouse anti-MCL-1 (Proteintech # 66026-1-Ig), rabbit anti-DRP-1 (Cell Signaling Technologies #8570S), rabbit anti-OPA1 (Cell Signaling Technologies #67589S), and control containing no primary antibody.

### Photoconversion experiments

Mitochondrial network connectivity and fusion was assayed using photo-conversion of mitochondria tagged with Mito-tdEos. Photo-conversion was performed on a Nikon Eclipse Ti inverted widefield microscope equipped with a 1.45 NA 100X Oil objective. Briefly, a stimulation region was closing down the field diaphragm and using the filter to shine 405 nm light for 6 seconds. Images for the converted (TxRed) and unconverted (FITC) were acquired before and after stimulation. The TxRed image before stimulation was used to subtract background from the post-stimulation images, followed by thresholding and automated measurement in Fiji (Schindelin et al., 2012). The initial converted area immediately after stimulation was used as a measure of connectivity, while the spread of the converted signal after 20 minutes was used as a measure of fusion/motility. The initial converted area (TxRed channel) was normalized to the total unconverted area (FITC channel) to account for any initial variation in the total mitochondrial area.

### Image acquisition

Super-resolution images for Figures 1 and 2 were acquired using a GE DeltaVision OMX microscope equipped with a 1.42 NA 60X Oil objective and a sCMOS camera. Super-resolution images for Figure 6 were acquired using a Nikon SIM microscope equipped with a 1.49 NA 100x Oil objective an Andor DU-897 EMCCD camera. Images for Figures S3, 4, and S5 were acquired on a Nikon Eclipse Ti inverted widefield microscope equipped with a 1.45 NA 100X Oil or 1.40 NA 60X Oil objective. Image processing and quantification was performed using Fiji. Measurement of cell number to assay cell death was performed on a Incucyte S3 live cell imaging system (Essen Bioscience) equipped with a 10X objective. Images for the PLA experiments were acquired on a Nikon spinning disk confocal microscope equipped with a 1.40 NA 60X Oil objective.

### Statistical Analysis

All experiments were performed with a minimum of 3 biological replicates. Statistical significance was determined by unpaired, two-tailed Student’s t-test or by one- or two-way ANOVA as appropriate for each experiment. GraphPad Prism v8.1.2 was used for all statistical analysis and data visualization.

Error bars in all bar graphs represent standard error of the mean or standard deviation as described for each figure, while Tukey plots were represented with boxes (with median, Q1, Q3 percentiles), whiskers (minimum and maximum values within 1.5 times interquartile range) and solid circles (outliers). No outliers were removed from the analyses.

For MEA experiments, means from triplicate biological replicates (each with three technical replicate wells) for each measurement were plotted and significance was determined by two-way ANOVA.

For PLA experiments, images were quantified using Fiji. Briefly, background noise levels were subtracted, and number of puncta per ROI was normalized to mitochondrial area. ROIs in at least 5 cells per condition were quantified in three independent experiments.

Quantification of actin organization was performed in a blinded fashion and percentages of each category are displayed. Cell viability measured using the Incucyte live cell imaging system was performed by automatic segmentation of nuclei in Fiji, followed by subtraction of dead cells as indicated by fragmented nuclei and rounded phenotype.

## Acknowledgements

We would like to thank Dr. Kevin Ess for providing access to the Axion Biosystems MEA analyzer, John Snow for providing critical technical support with the Axion analyzer, Bryan Millis for providing expertise with high resolution microscopy, and Stellan Riffle for technical support. This work was supported by 1R35 GM128915-01 NIGMS (to VG), 4R00CA178190 NCI (to VG), 1R21CA227483-01A1 NCI (to VG), 19PRE34380515 AHA (to MLR), 18PRE33960551 (to NT), and R35 GM125028-01 (to DTB). The Vanderbilt Cell Imaging Shared Resource is supported by NIH grants 1S10OD012324-01 and 1S10OD021630-01.

The authors declare no competing financial interests.

## Author contributions

V. Gama, M. Rasmussen and N. Taneja conceived the study, designed experiments, interpreted data, and wrote the manuscript. M. Rasmussen and N. Taneja designed and carried out all the cell biology experiments, with input from D. Burnette, A. Neininger, L. Wang, and L. Shi. V. Gama designed and supervised the project. The manuscript was prepared by M. Rasmussen and V. Gama, and revised by N. Taneja and D. Burnette. D. Burnette and B. Knollmann provided vital reagents and critical expertise.

The authors declare no competing interests.

