## Supplementary figures and images for "MCL-1 inhibition by selective BH3 mimetics disrupts mitochondrial dynamics in iPSC-derived cardiomyocytes"

### Supplemental Figure 1

Supplementary Figure 1

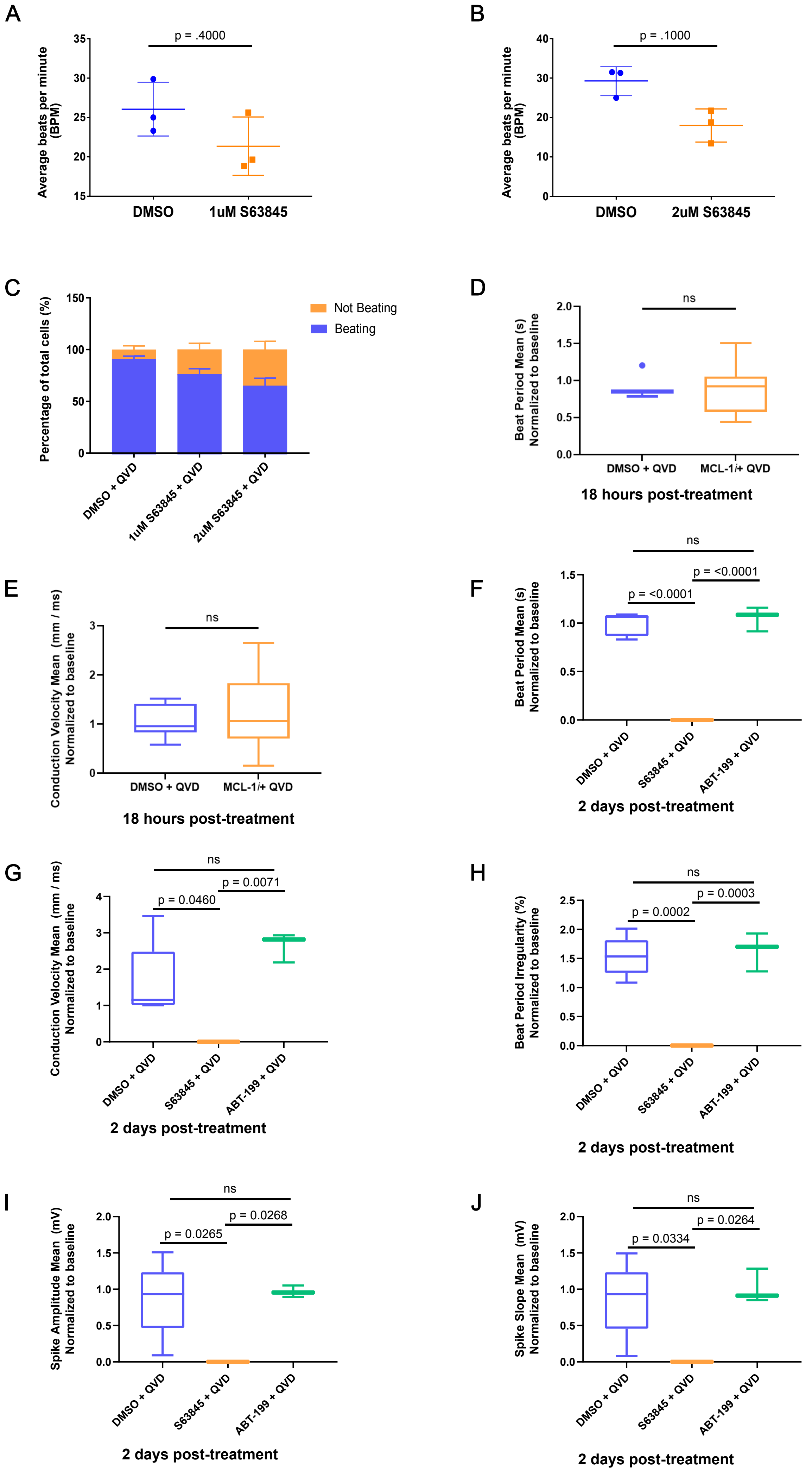

### Supplemental Figure 2

A

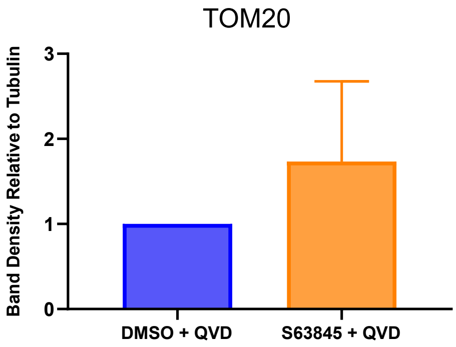


B

**Secondary PLA probes only**

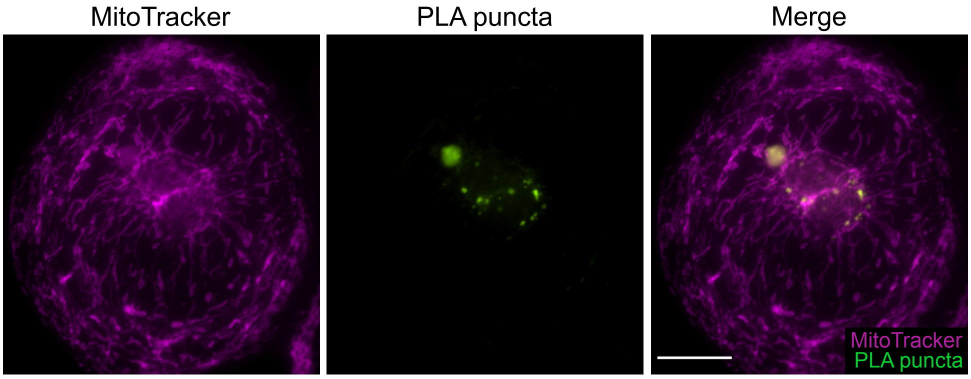

### Supplemental Figure 3

#### Supplementary Figure 3

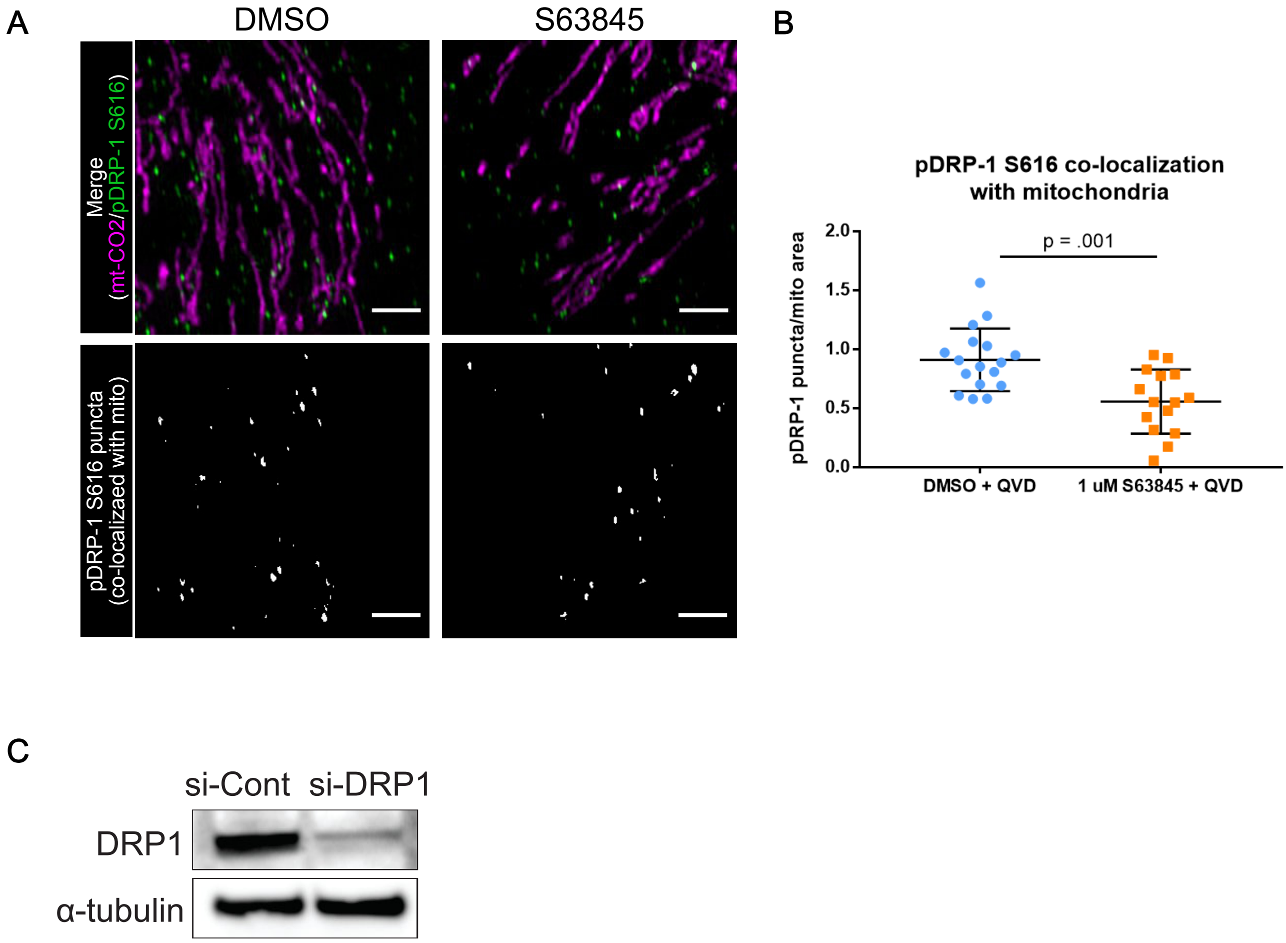

### Supplemental Figure 4

Supplementary Figure 4

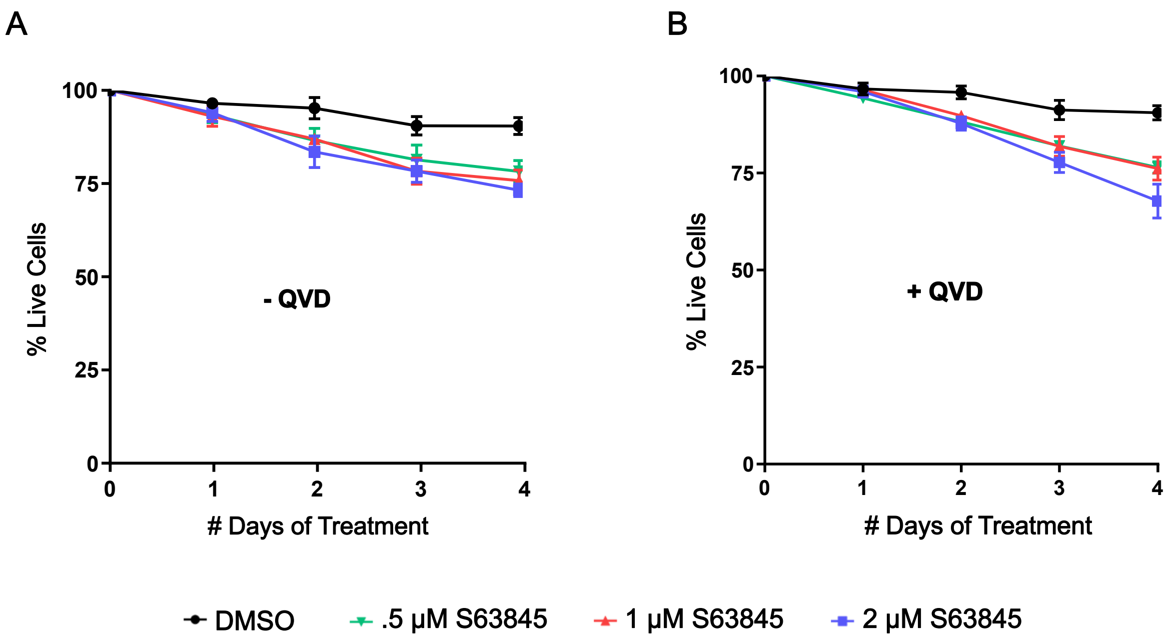

### Supplemental Figure 5

Supplementary Figure 5

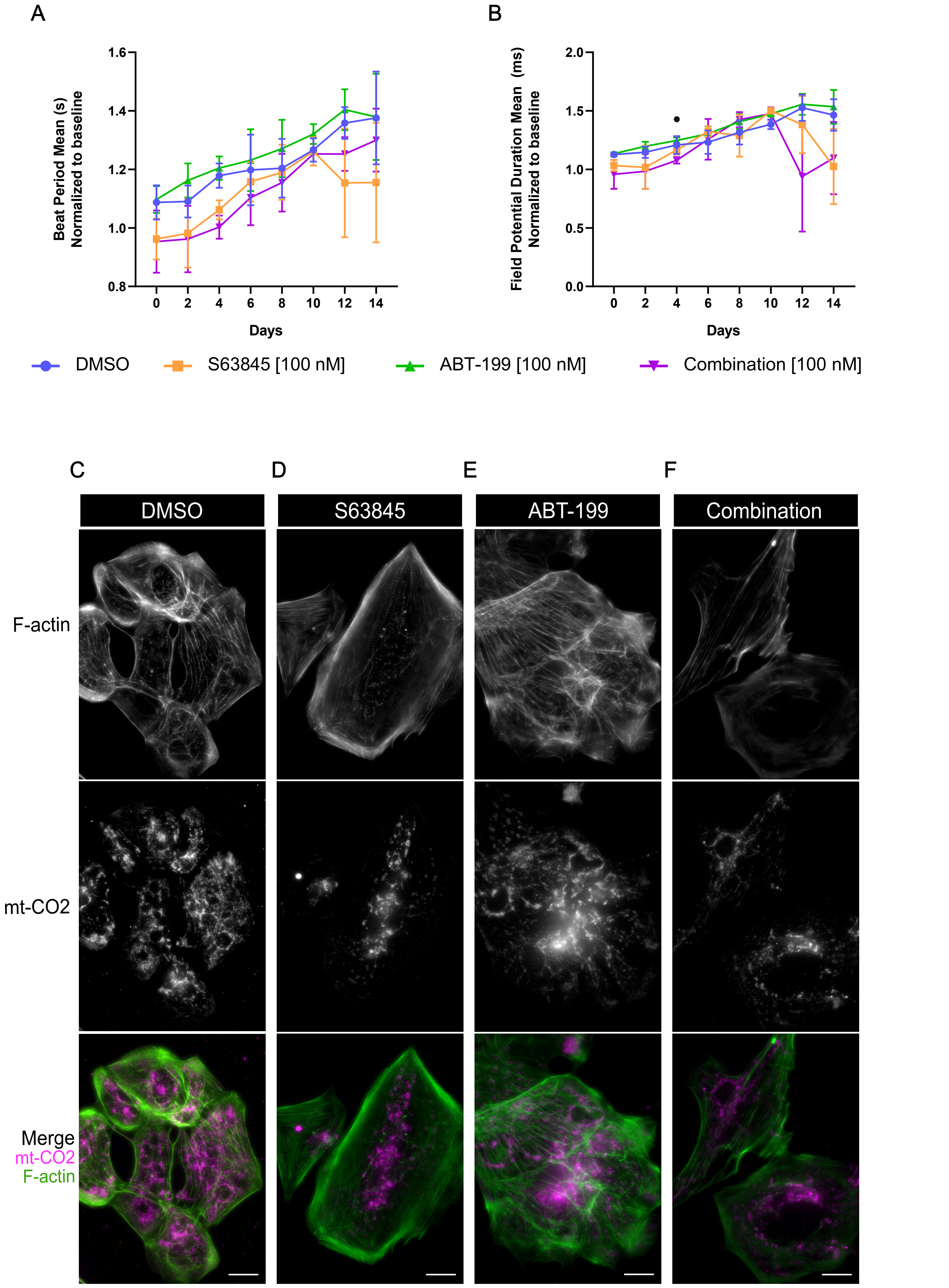
