## Supplemental Figure legends for "MCL-1 inhibition by selective BH3 mimetics disrupts mitochondrial dynamics in iPSC-derived cardiomyocytes"

**Figure S1: Corresponding to Figure 2.** (A-B) hiPSC-CMs were plated on a CytoView MEA plate (Axion Biosystems) and treated with either vehicle (DMSO) or .5 μM MCL-1*i* (S63845) and QVD. Live-cell activity was recorded for 5 minutes and results were normalized to baseline recordings for each well. Beat period (A) and conduction velocity mean (B) are not significantly different between control and treated cells at 18 hours-post-treatment. (C-G) hiPSC-CMs were plated as in Figure S1A-B and treated with either vehicle (DMSO), .5 μM MCL-1*i* (S63845), or .5 μM ABT-199 and QVD. After 2 days, ABT-199-treated cells display normal beat period mean (C), conduction velocity mean (D), beat period irregularity (E), spike amplitude mean (F) and spike slope mean (G). By the 2-day time point, all MCL-1*i*-treated cells were quiescent and no longer displayed any measurable activity (C-G). (H) hiPSC-CMs were treated every other day for 4 days and beat rate was recorded using live-cell phase contrast imaging. Percentages of beating cells were compared to non-beating cells for increasing doses of MCL-1*i* (Chi-square test p = <0.0001). Average beats per minute (BPM) are shown for 1 μM MCL-1*i* (I) and 2 μM MCL-1*i* (J) in combination with QVD. Error bars indicate ±SD for bar graphs, boxplots are shown as Tukey graphs.

**Figure S2: Corresponding to Figure 3.** (A) Quantification for TOM20 band density from the experiment shown in Figure 3C. Error bars indicate ±SD. (B) Representative image of secondary PLA probe control sample. The area around the nucleus was avoided for all quantification purposes.

**Figure S3: Corresponding to Figure 4.** (A) hiPSC-CMs were treated every other day for 4 days with DMSO or 1 μM MCL-1*i* and QVD, then fixed for immunofluorescence. Samples were stained with antibodies against mt-CO2 (mitochondria) and pDRP-1 S616 (active DRP-1), and images were quantified for number of pDRP-1 S616 puncta co-localized with mitochondria (B). Representative images are shown. Scale: 2 μM. Error bars denote ±SEM and experiment was performed in 3 independent replicates. (C) Representative Western blot showing knockdown of DRP-1 compared to control siRNA in the experiment shown in Figure 4E-F.

**Figure S4: Corresponding to Figure 5.** (A) hiPSC-CMs were treated with S63845 at the indicated concentrations and either vehicle DMSO (F) or with QVD (G). Cell survival was measured using an Incucyte incubator. Representative images from each day of treatment were quantified and percentages of live cells normalized to total cells are shown. Error bars denote ±SEM and experiment was performed in 4 independent replicates.

**Figure S5: Corresponding to Figure 6.** Chronic inhibition of MCL-1, but not BCL-2, results in cardiac activity defects. hiPSC-CMs were treated every 2 days with DMSO (blue), 100 nM MCL-1*i* (S63845 - orange), 100 nM BCL-2*i* (ABT-199 - green), or both inhibitors (magenta) for 14 days. MEA plate was recorded 2 hours-post-treatment for 5 minutes and results were normalized to baseline recording for each respective well. Results of recordings for beat period mean (A) and field potential duration mean (B) are shown. Error bars show ±SEM. P-value is shown as ● = ABT-199 vs. Combination. One symbol indicates p = <0.05. (F-I) Mitochondria and F-actin were imaged at the end of the treatment paradigm in Figure 6A-E. Representative SIM images are shown of cells treated with DMSO (F), 100 nM MCL-1*i* (S63845) (G), 100 nM BCL-2*i* (ABT-199) (H), and 100 nM MCL-1*i* + 100 nM BCL-2*i* (Combination) (I). Scale: 10 μm.
